# Men in menopause? Experimental verification of the mate choice theory with *Drosophila melanogaster* shows both sexes can undergo menopause

**DOI:** 10.1101/2022.04.18.488714

**Authors:** Divya Purohith, Mitali Chaudhary, Alyssa Gomes, Nina Rajapakse, Aditi Das, Neha Dhanvanthry, Michelle Brown, Manan Mukherjee, Rama S. Singh

## Abstract

Various hypotheses regarding the origin of menopause have been proposed, and although the kin-selection-based theory appears promising, it involves population genetic processes that are insufficient to compensate for loss of fitness. The grandmother hypothesis and its variation the live long hypothesis are untenable; the former requires “climbing a steep fitness hill”, as grandmothers share only 25% of their genes with their grandchildren, compared to 50% with their direct offspring, while the latter proposes a prolongation of the post-menopausal lifespan through selection, which is impossible in a population of non-reproducing females. The mate choice theory explains menopause as the result of asymmetric mating involving younger females and older males that leads to an accumulation of infertility mutations and the evolution of menopause in older females. In this study, we investigated the mate choice theory using an infertility mutation accumulation experiment with *Drosophila melanogaster* that involved mating between individuals of different age groups. After 70 generations of asymmetric mating, the results showed that younger females who were paired with older males showed declining fertility in old age. The same trend was noted with younger males when mated with older females; the fertility of the males declined in old age. These results support the mate choice theory and indicate that menopause is not a life history trait of females but of the sex of the younger mate. Mate choice theory treats the evolution of menopause and post-menopausal lifespan as independent traits that are driven by the mate choices exercised by older males. Menopause may be an atypical process because the evolutionary mechanism (age-restricted asymmetric mating) involved is rarely observed.

## Introduction

Human menopause is defined as a complete cessation of the menstrual cycle that equates to the loss of fertility and is usually characterized one year after the final menses of a female [1]. This phase is often called “post-reproductive life,” signifying that reproductive senescence predates biological senescence [2]. Natural selection should operate against both the early termination of reproduction and survival beyond reproductive senescence [3]. This post-reproductive life trait was thought to be unique to humans [4]; however, it has been shown to occur in several species of toothed whale [5,6] and, less commonly, in captive chimpanzees [7,8]. Other mammals continue to reproduce until death [9]. This evolutionarily counterintuitive [10] cessation of reproduction in human females occurs at approximately 50 years of age and is preceded by years of diminishing fertility [11]. As evolution favors reproduction, the decline of reproduction in older females is an evolutionary puzzle [12].

Two perspectives on human menopause are considered: a medical perspective that focuses on the health of post-menopausal females [13] and an evolutionary perspective that focuses on the origin and evolution of menopause [12]. Under the medical perspective, cessation of reproduction following menopause results from the continuous and lifetime loss of a limited supply of follicles from the ovaries that are maximized at birth [13]. Although follicular loss occurs through ovulation, the majority of the loss is likely from the degeneration of non-ovulating follicles (programmed atresia) with ovulation accounting for only a small portion [14]. In addition, a decrease in oocytes with age might occur due to genetic defects in these cells over time as a result of unrepaired meiotic DNA damage or a breakdown in DNA repair mechanisms [15–17].

The evolutionary perspective can be adaptive or non-adaptive (see Table 1). Under the non-adaptive hypotheses, menopause is an epiphenomenon: an indirect result of selection due to antagonistic pleiotropy that favors early reproduction; alternatively, menopause may be a by-product of the increase in lifespan (the “lifespan artifact hypothesis”) [18,19]. The increase in lifespan is believed to be based on intrinsic genetic or social and cultural factors [18,20–23]. While the non-adaptive models could account for the evolution of menopause in females [18,24], they do not explain the retention of fertility in males [12].

**Table 1.** Menopause hypotheses and sources, modified from Morton et al. (2013)

| **Hypothesis** | **Description** |
| --- | --- |
| Follicle-depletion hypothesis | Females have a fixed number of follicles and menopause ensues when the follicles are depleted [27,28]. |
| Lifespan artifact hypothesis | Menopause is an automatic process that occurs when human longevity is increased [19,29]. |
| Grandmother hypothesis | Menopause allows grandmothers to contribute to the survival of their grandchildren, thus increasing their inclusive fitness [25]. |
| Reproduction-cost hypothesis | Investment in reproduction is higher in females than males, leading to physiological deterioration and susceptibility to infertility [30]. |
| Mother hypothesis | (An adaptive version of the reproduction-cost hypothesis) Aging mothers increase the survival probability of their children; menopause also prevents fertilization of nonviable ova [31]. |
| Patriarch hypothesis | The origin of menopause allowed males to mate with younger females, thus increasing the longevity of both sexes, as well as the status of males in society [32]. |
| Senescence hypothesis | Menopause is the natural effect of aging. Reproductive aging precedes somatic aging [33]. |
| Absent father hypothesis | Reduced paternal investment and increased maternal age induce menopause. This complements the grandmother hypothesis [34]. |
| Evolutionary trade-off | Menopause is a trade-off between the future reproduction of a female and the enhanced survival of her existing offspring [35]. |
| Reproduction-conflict | Menopause is the evolutionary outcome of resource-based competition between generations (i.e., between a grandmother and her daughters-in-law, who are non-blood relations); on the basis of genetic relatedness, fitness can be optimized if non-blood relations reproduce with the help of grandmothers [21,29]. |
| Mate choice theory | A bias in mating behavior, where only younger females are allowed to mate, allows late-onset fertility-reducing mutations to accumulate and fix in the population, ultimately leading to menopause [12]. |
| Inclusive mate choice/triumvirate hypothesis approach | A combination of mate choice theory, lifespan extension, and grandmother assistance in rearing grandchildren [26]. |

Under adaptive hypotheses such as the grandmother theory, menopause is considered an adaptive trait: the direct result of favorable selection of post-reproductive lifespan that allows menopausal females to recuperate their loss of fitness through “inclusive fitness” by investing in their grandchildren [25]. The “live long” hypothesis is a variation of the grandmother hypothesis that suggests that post-menopausal females have extended their lifespan beyond their reproductive years [6]. Kin-selection-based hypotheses may appear promising; however, the principles of population genetics deem these theories unfeasible. For a menopausal female, recuperating loss of fitness through “grandparenting” is an “uphill battle,” as grandmothers share only 25% of their genes with their grandchildren as compared to 50% with their offspring.

Therefore, for every loss of a child, a grandmother would be required to raise two additional grandchildren. Furthermore, irrespective of gain through inclusive fitness, these theories do not explain how menopause or infertility evolved, regardless of whether it developed before or after the lifespan extension [26].

One non-kin-selection and non-adaptive mate choice hypothesis postulated that a specific change in mating behavior (the preference of males for younger females) could have led to the evolution of menopause [12]. Under such a mating system, selection would be relaxed in older females who were deprived of reproduction, and late-onset fertility-reducing mutations would be rendered neutral and accumulate, leading to the cessation of reproduction and evolution of menopause [12]. Although this theory is propitious and applicable to both sexes and all sexually reproducing organisms, it has not been examined experimentally.

We investigated the mate choice theory using *Drosophila melanogaster* and examined the role of preferential, age-based asymmetric mating (henceforth, “asymmetric mating”) in the origin and evolution of fertility-reducing mutations leading to menopause. We examined the effect of asymmetric mating in both sexes by mating younger females with older males and younger males with older females. The rationale behind the experiment was that the deprivation of reproduction in older individuals would result in the accumulation of sex-specific infertility mutations and sterility in older flies, irrespective of sex.

Our study involved an asymmetric mating experiment that covered 70 generations. The results confirmed those predicted by the mate choice theory [12]: female flies used as young mates (henceforth “young-female population”) and males used as young mates (henceforth “young-male population”) became infertile in old age. The results indicate that menopause is not a life-history characteristic that is specific to females but occurs in the sex which is mated at a young age. In both experiments, the younger sex developed “menopause” while the older sex remained fertile.

## Results

The mutation accumulation experiment was designed to cover 100 generations; however, the COVID-19 pandemic forced the project to be discontinued after the 70^th^ generation. Fitness observations were made at three points for Generations 20, 24, and 70 and included fecundity, fertility, longevity, ovariole number, testis length, and testis width. We also examined the lifespan of Generation 24 to determine if it was affected by the experimental treatment, as lifespan and infertility are considered to be linked in discussions of menopause.

### Measurement of fertility: Generations 20 and 24

#### Accumulation of deleterious mutations negatively affected fertility

Although we had not expected to observe the effects of new infertility mutations in the early generations, measurements were taken for comparative purposes and to determine whether asymmetric mating had any initial effect on fecundity and fertility, independent of mutations.

The results are summarized as follows: (1) The experimental lines laid slightly fewer eggs than the control group on Day 3 post-mating, after which egg numbers declined linearly until Day 16 (Fig. 1A); (2) a large reduction in the number of offspring produced in the experimental lines was observed (Fig. 1B); (3) the egg fertility or “hatchability” (ratio of offspring/eggs) test showed a significant reduction in egg hatchability between the control and experimental lines (Fig. 1C); and (4) 18 days of measurement revealed that a complete loss of fertility in the experimental lines occurred long before female mortality. In contrast, the control females reproduced throughout their lifetime, although at a lower rate (Fig. 1D).

**Fig. 1.**
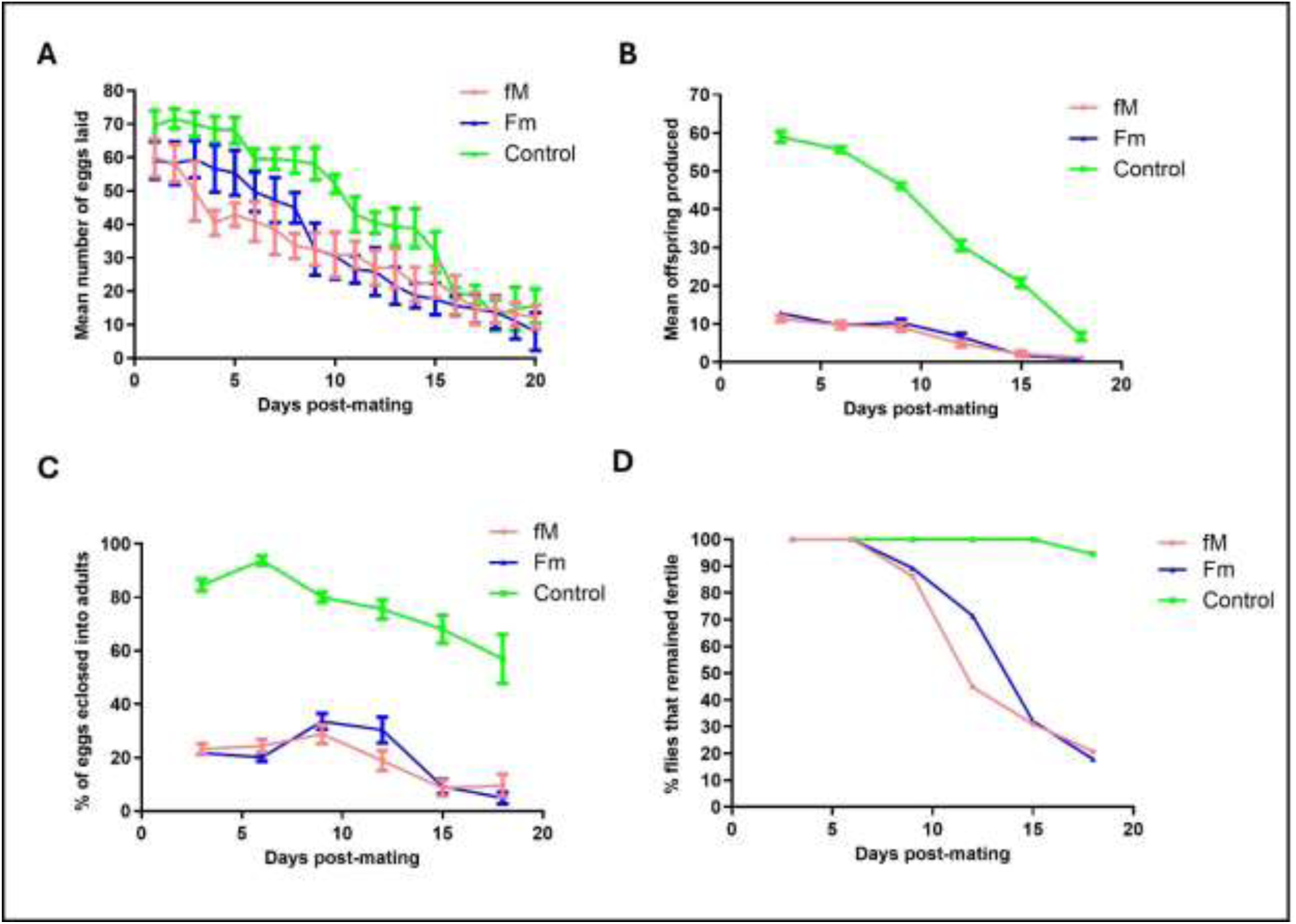
Fecundity and fertility curves for the control and experimental (fM and Fm) populations. (A) Mean number of eggs laid per day by females from control and experimental lines (fM, Fm), (B) mean number of offspring produced per day, (C) percentage of eggs eclosed per day, and (D) percentage of flies remaining fertile as a function of age, measured every third day. Bars represent the standard error of the mean.

The most likely cause of the initial loss of fecundity and fertility in the experimental populations on Day 3 was the developmental immaturity of the young flies used in the mating experiments. This initial loss may or may not be reversed in later generations, but the faster rate of complete loss of fertility in the experimental lines compared to the control after Day 3 may indicate the contribution of mutation accumulation.

The reduction in the fecundity and fertility of experimental lines on Day 3 was repeated in Generation 24 using only one experimental population (fM) and the control. We also measured fitness-associated traits (egg chamber number, ovariole number, testis length, and testis width), survivorship, and lifespan (for both sexes). The results are presented in Fig. 2 and show a pattern similar to that observed with Generation 20 (Fig. 1). The number of eggs laid, number of offspring produced, and hatchability of the eggs were lower in the experimental (fM) population when compared to the control (Fig. 2A, B, C). The analysis of variance (ANOVA) showed a significant difference between the experimental flies and the control (p < 0.0001). Similar to the results for Generation 20, the complete loss of fertility occurred faster in the experimental population (fM) than in the control, with 22% of the experimental (fM) flies and 88% of the control flies remaining fertile by Day 18 (Fig. 2D).

**Fig. 2.**
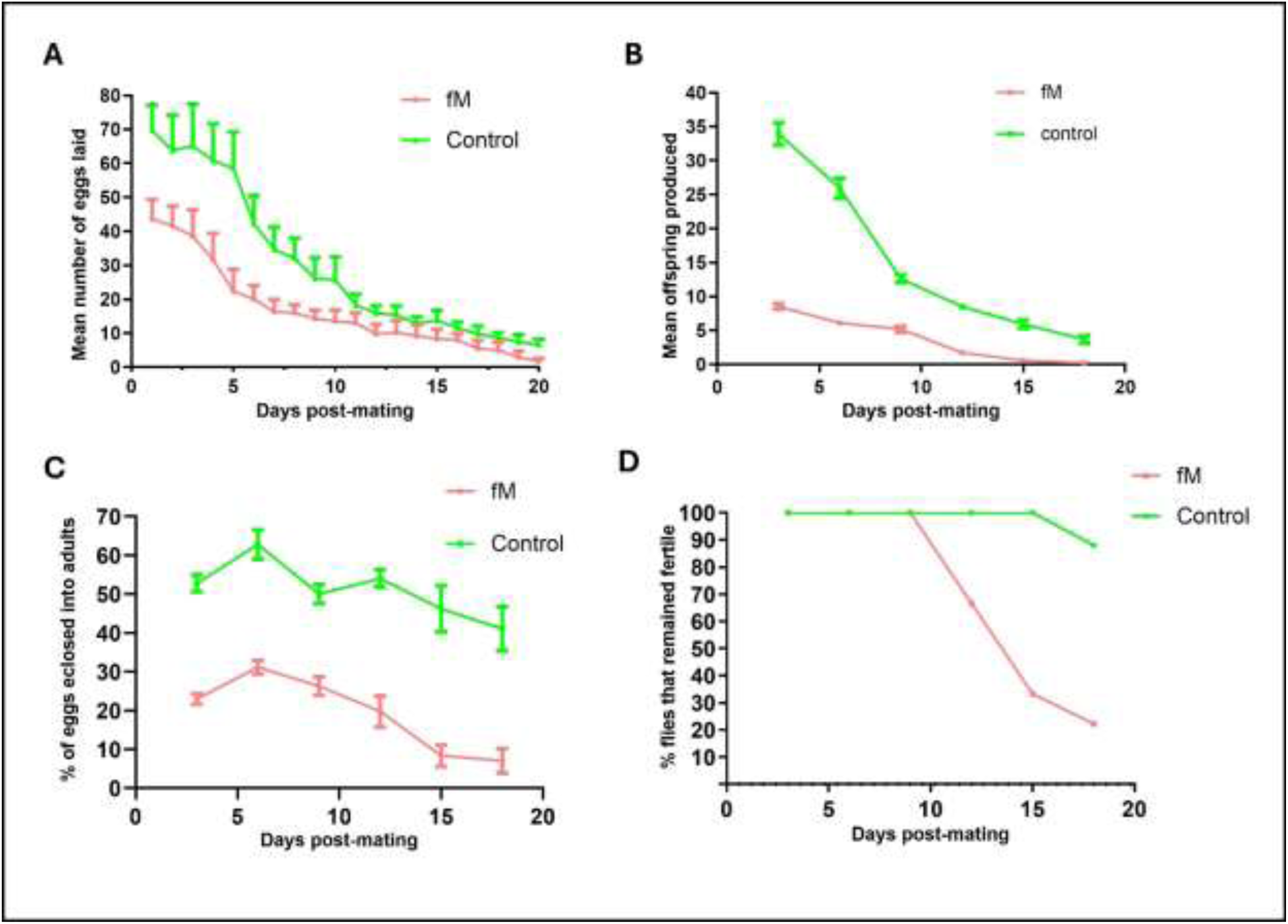
Fecundity and fertility curves of the control and young female (fM) experimental populations. (A) The mean number of eggs laid per day, (B) mean number of offspring eclosed per day, (C) percentage of eggs eclosed, and (D) percentage of female flies remaining fertile as a function of age. Bars in all figures represent the standard error of the mean.

We used Weibull models (see Material and Methods section) to analyze the data for the flies that remained fertile with age and an analysis of variance (ANOVA) to determine the best-fit model. The ANOVA of Generation 20 flies that remained fertile in experimental population fM (Fig. 1D) indicated that Model 2 (which allows for different parameters for each group) provided a significantly better fit of the data compared to Model 1 (which assumes the same parameters for both groups, i.e., the Null Model). The very low p-value (2.182 · 10^−06^) strongly suggests that the differences in parameters between the groups are statistically significant and sufficient evidence exists to reject the Null Model (Fig. 3 A, B). Similar results were obtained for experimental population Fm in Generation 20 (Fig. 1D), whereby Model 2 provided a significantly better fit to the data than Model 1 and the very low p-value (1.774 · 10^−07^) signifies a rejection of the Null Model (Fig. 3 C, D).

**Fig. 3.**
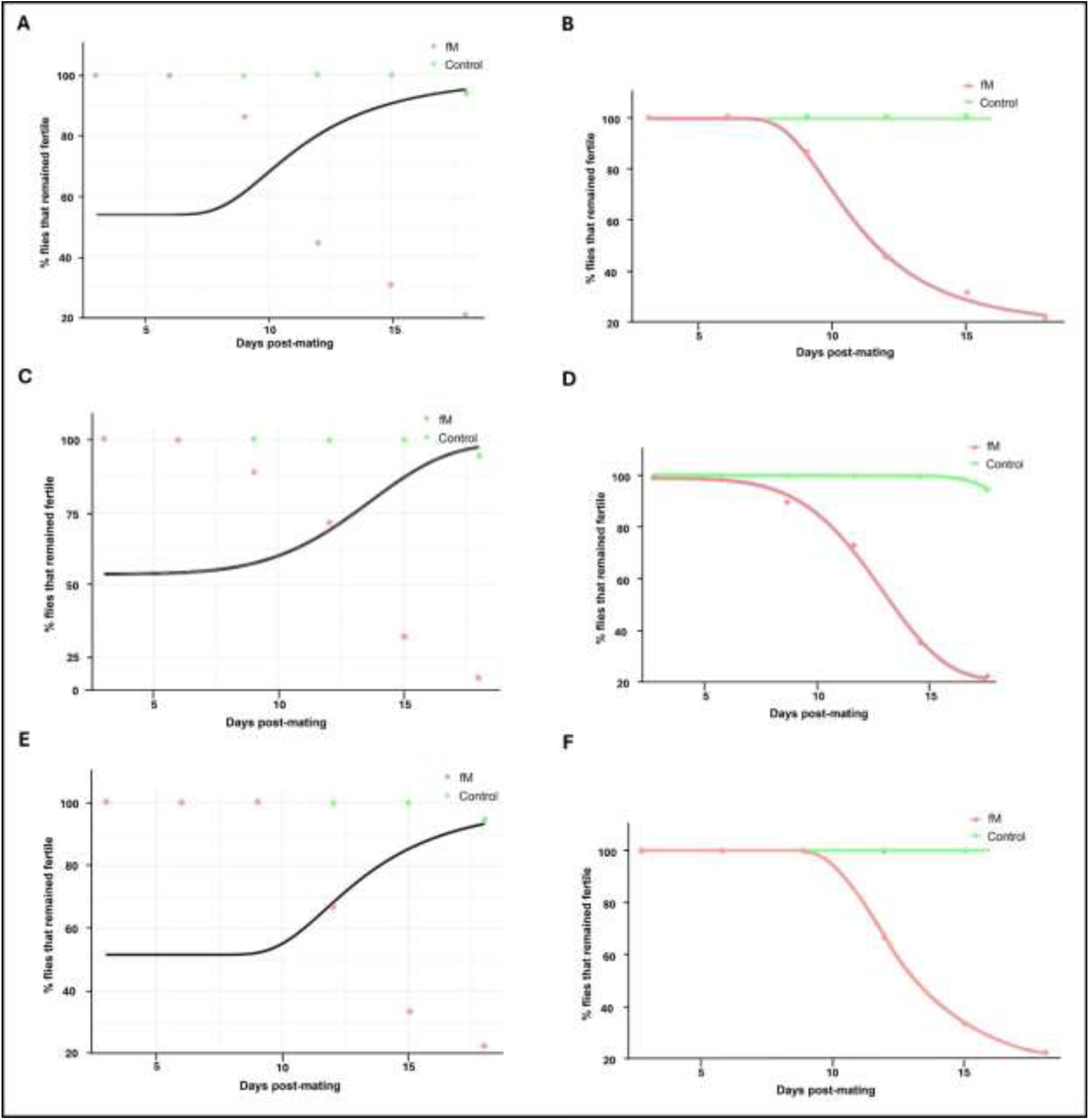
(A) The null model predicting the proportion of eggs that hatched over time, including both the control and experimental groups (fM) of Generation 20 (Mitali’s data), and the alternative model (B) where the control group (in orange) and experimental group (in cyan) (fM) are represented with independent parameters. (C) The null model predicting the proportion of flies remaining fertile over time, including both the control and experimental groups (Fm) of Generation 20 (Mitali’s data), and the alternative model (D) where the control group and experimental group (Fm) are each represented with independent parameters. (E) The null model predicting the proportion of eggs that hatched over time, including both the control and experimental groups (fM) of Generation 24 (Divya’s data), and the alternative model (F) where the control group (in orange) and experimental group (in cyan) (fM) are represented with independent parameters.

The ANOVA of flies remaining fertile in experimental population fM in Generation 24 also indicated that Model 2 provided a significantly better fit to the data than Model 1. The very low p-value (06 · 10^−07^) strongly suggests that the differences in parameters between the groups are statistically significant and the Null Model should be rejected (Fig. 3 E, F). These results reveal that the (decline in fertility was accelerated for both experimental populations (fM and Fm) when compared to the control.

### Effect on fitness-related traits

#### Survivorship and lifespan remained unaffected

The average expected lifespans for *D. melanogaster* females and males are 53 and 33 days, respectively [36,37]. Virgin females were observed to live longer than virgin males in all populations. Three populations were identified as outliers and removed from the analysis to reduce bias: one young-male population (2f) and two young-female populations (1f and 2m). Survival curves showed that the males had similar decreases in their lifespans over time, and the Chi-square test (p = 0.409) revealed that the difference between the control and the experimental male populations was not significant. Similar results were found for the female flies, whereby the Chi-square test (p = 0.34) indicated a non-significant difference between the control and experimental populations. Since one of the parents used in each experimental line was 20 days old (male in fM, female in Fm), we did not expect to see any effect on survivorship and lifespan, as any effect of mutation accumulation affecting survival in the younger sex would have been counteracted by the genes of the older parent.

#### Egg chamber and ovariole numbers were reduced in the young-female population (fM)

The mature eggs (stage-14 egg chambers) were counted every third day post-eclosion (Fig. 4A). The number of egg chambers was similar in the experimental (Fm) and control populations, decreasing from 10-12 on Day 3 to 3-4 on Day 30. As expected, the fM population produced the fewest number of stage-14 egg chambers throughout the observation period. The one-way ANOVA (p = 0.0034) rejected the null hypothesis of no difference among populations, while the two-tailed *t*-test showed a significant difference between the number of mature egg chambers in the fM population and the number in the control group (control vs. fM; p = 0.0007). In addition, a significant difference was observed between the two experimental populations (fM vs. Fm; p = 0.014).

**Fig. 4.**
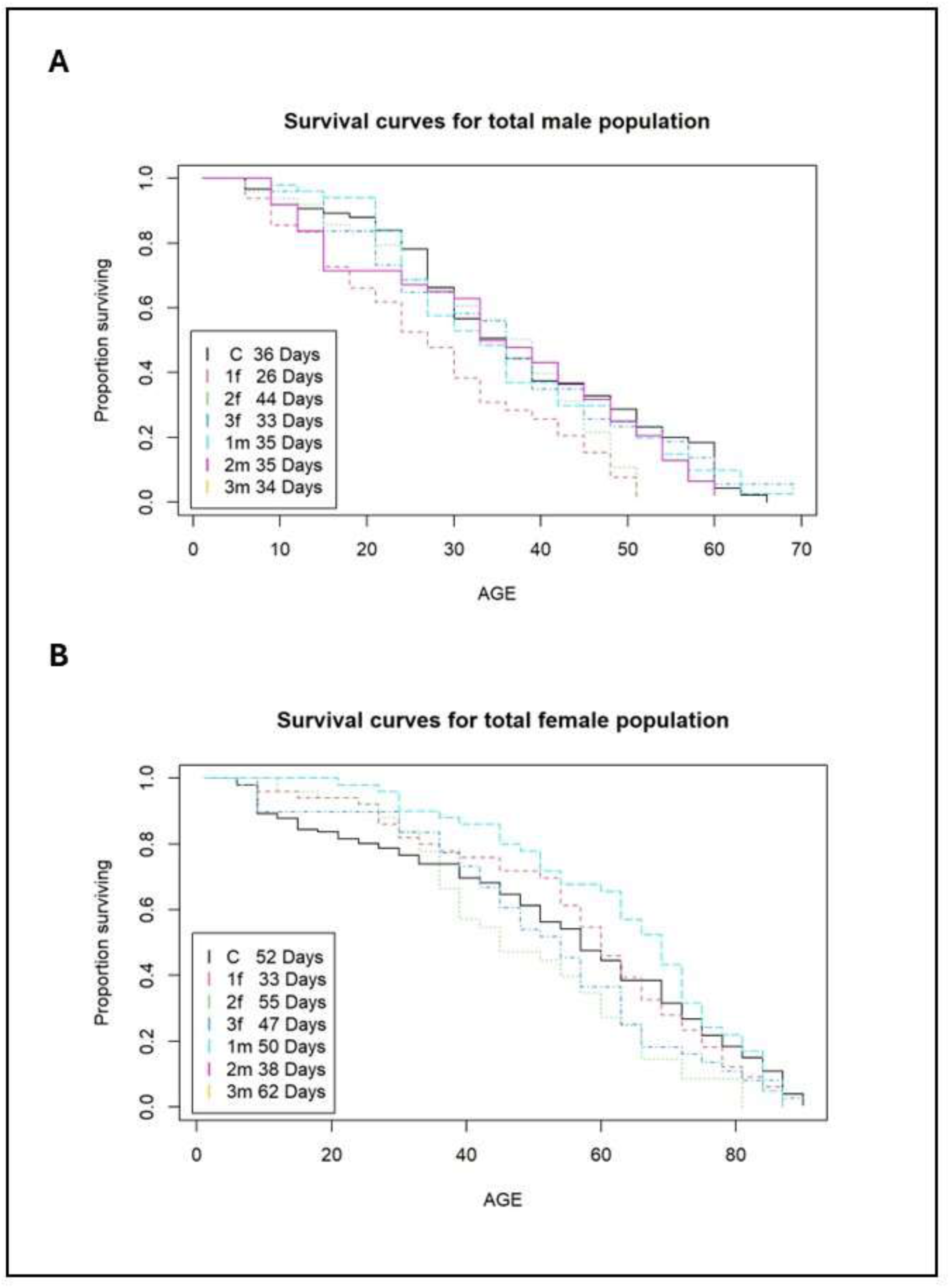
Survival curves for the total number of males and females from the pooled control and two experimental populations. Expected lifespans are included within the figures. The y-axis is the proportion of flies surviving at each interval, and the x-axis is the age of the flies (days). The three replicates of the young-female (fM) population are denoted as 1f, 2f, and 3f; the three replicates of the young-male (Fm) population are designated as 1m, 2m, and 3m; and the control is identified as C.

The number of ovarioles per female is represented in this study by the average number in the left and right ovaries. This number fluctuated between 14 and 18 until Day 21 when a sharp decline occurred in all lines (both experimental and control). The one-way ANOVA (p = 0.0001) rejected the null hypothesis of no difference among populations, and the two-tailed *t*-test showed a significant difference in the ovariole count between the fM experimental population and the control (p < 0.0001). Reductions in ovariole numbers and egg counts occurred in parallel in the young-female experimental population (fM).

#### Testis width, but not testis length, declined in the young-male population (Fm)

The testis length and width from ten individuals from each of the three replicates of the experimental and control populations were measured as a proxy of male fertility (Figs. 5,6).

**Fig. 5.**
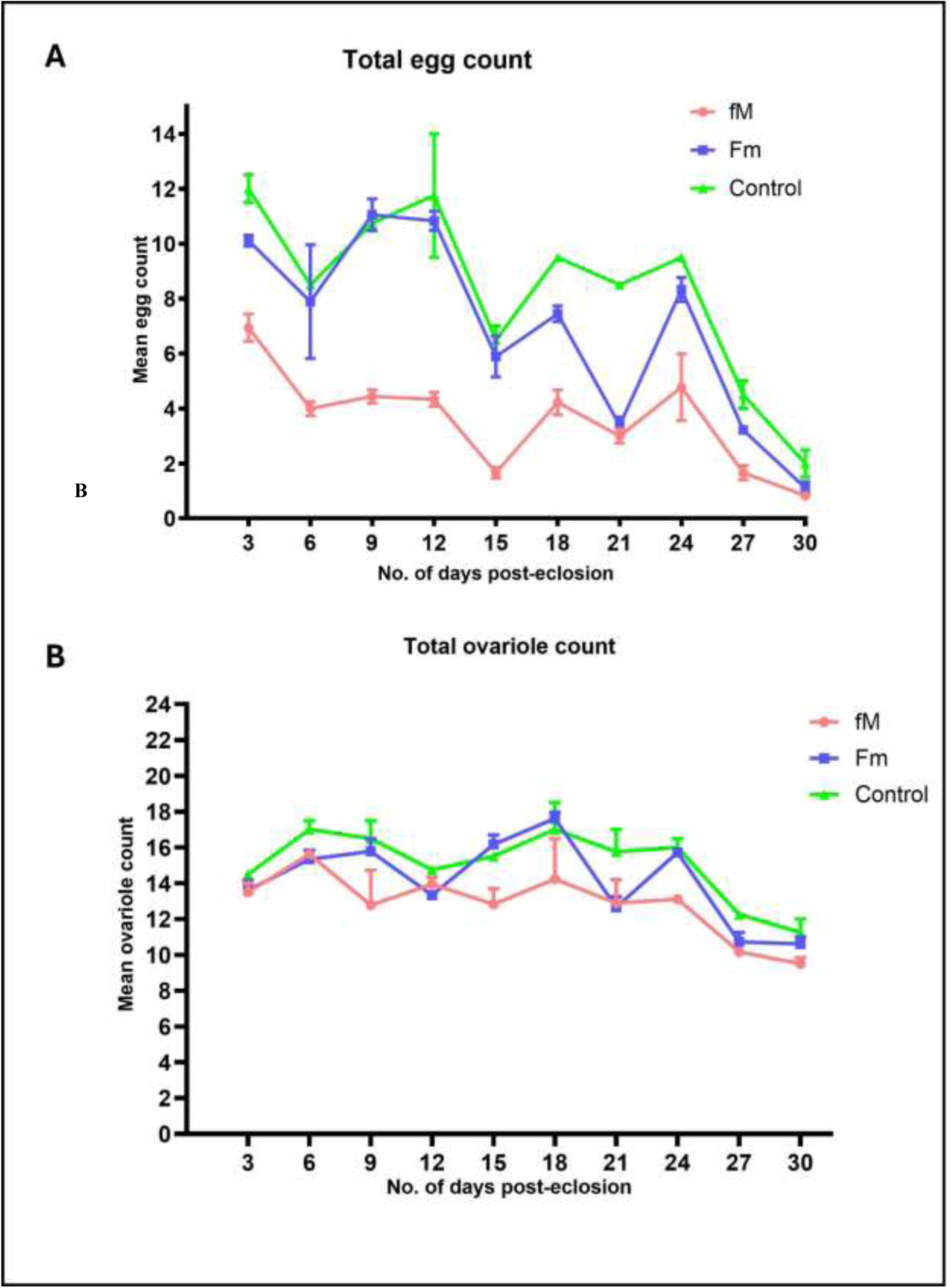
A Plot of stage-14 egg chamber and ovariole counts for the control and two experimental populations. (A) Mean number of stage-14 egg chambers counted per day, and (B) mean number of ovarioles counted per day. Observations were made every third day. Bars in the graph represent the standard error of the mean.

**Fig. 6.**
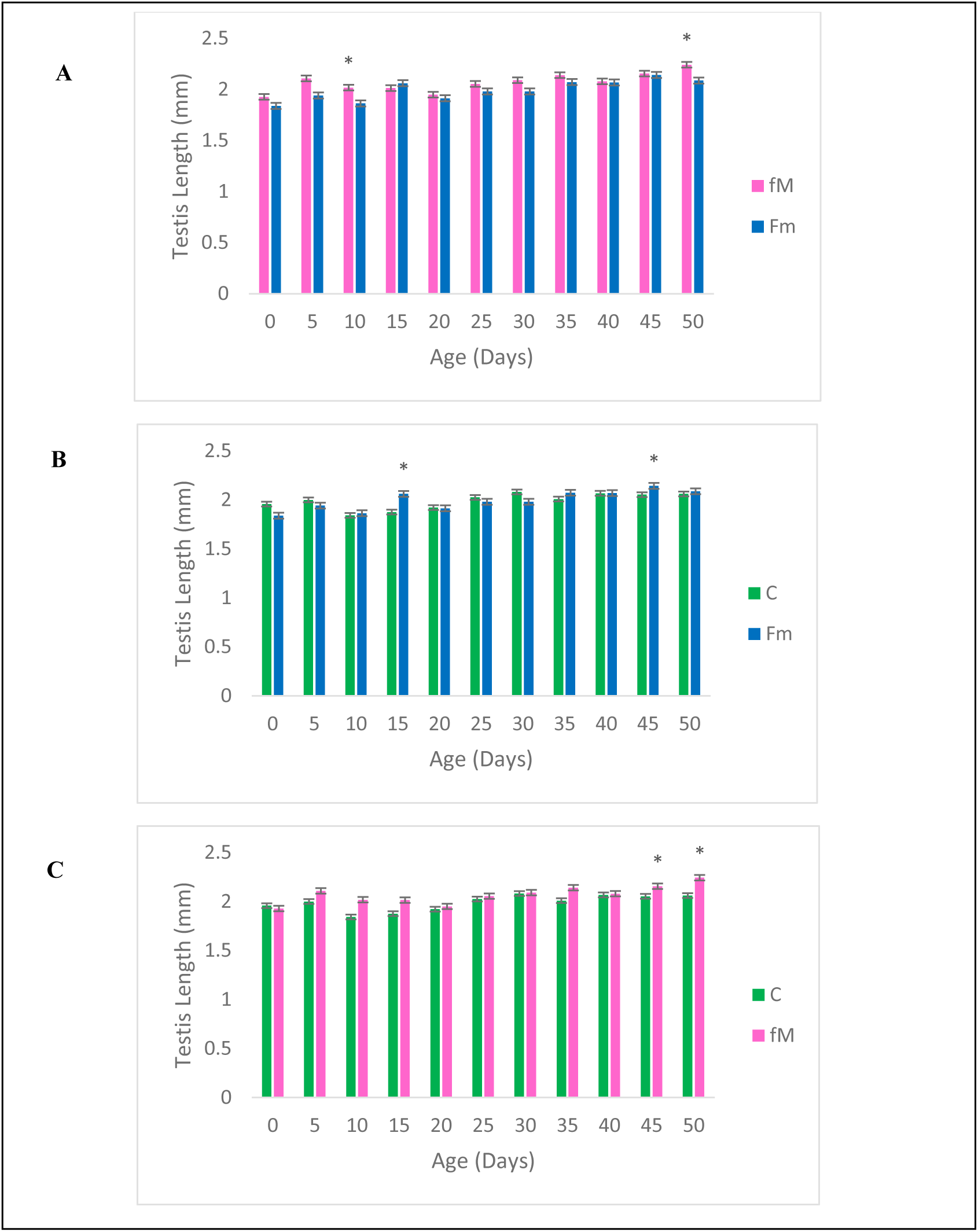
Pooled (over three replicates) mean testis length for Fm, fM, and control (C) populations. Comparison of mean testis length of fM vs. Fm (A), Fm vs. control (B), and fM vs. control (C). *p <0.05. Error bars denote the standard error of the mean.

**Fig. 7.**
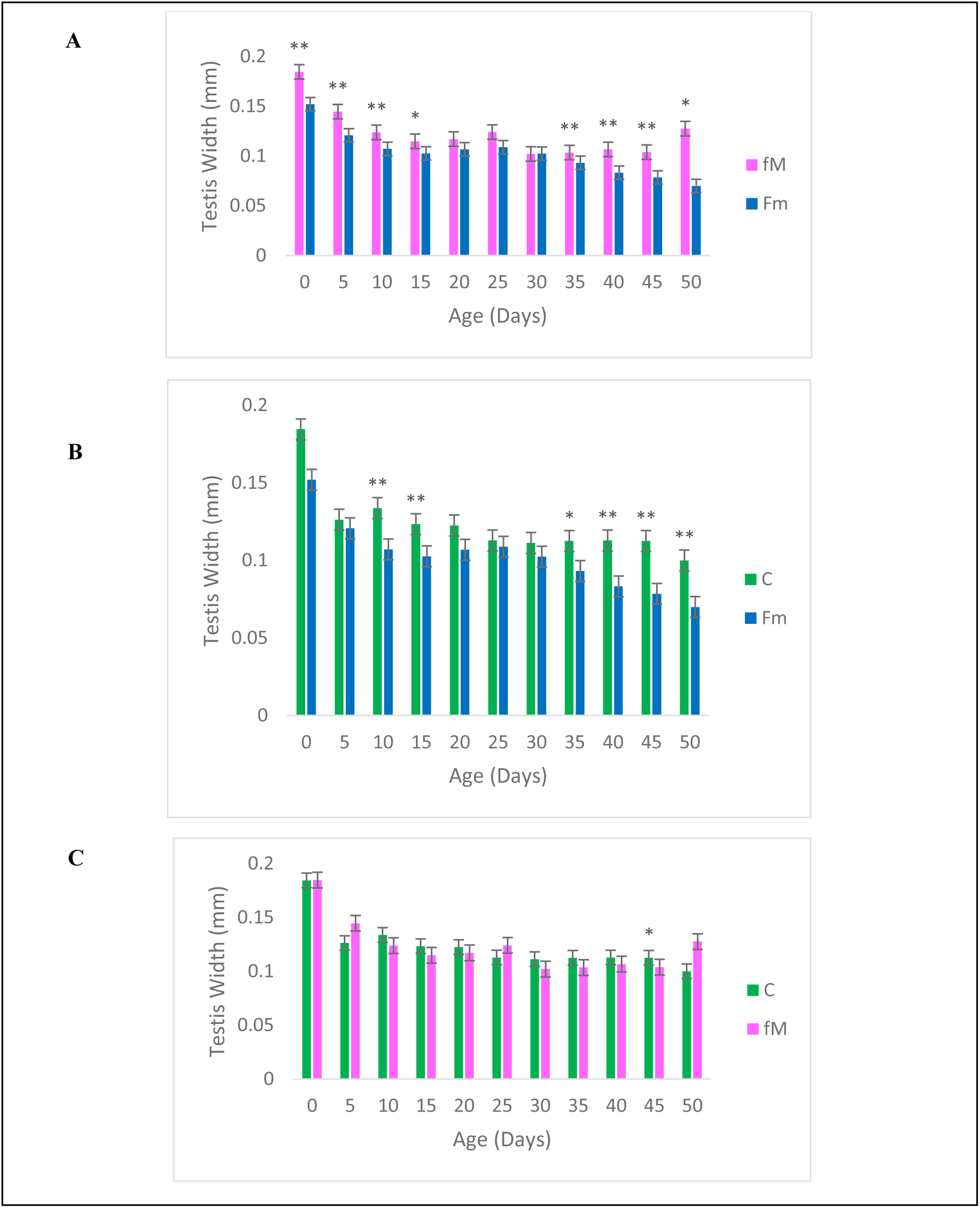
Pooled (over three replicates) mean testis width for Fm, fM, and control (C) populations. *p <0.05, **p <0.02. Error bars denote the standard error of the mean. Comparison of mean testis width of fM and Fm (A), Fm and control (B), and fM and control (C).

Testis length showed no significant variation during the observation period (50 days); however, the testis from the Fm population was narrower than that of the control and the fM, especially after Day 25 (Fig. 6 A, B). The two-tailed *t*-test showed mean testis width was consistently and significantly lower in the Fm group than the control and fM populations during the late life period (Days 25–50) (Fig. 6A, B). Male fertility is expected to be more sensitive to mutational variation than female fertility and the decline in late-age testis width may result from fertility-related mutational deterioration.

### Measurement of fertility: Generation 70

#### Fertility declined in both the young-female (fM) and young-male (Fm) experimental populations

Flies from Generation 70 were used for the second round of fitness measurements. The experiment was discontinued after this stage due to the COVID-19 pandemic. The same components of fertility that were measured in Generations 20 and 24 were obtained for Generation 70, and the results are shown in Figs. 8 and 9. The egg hatchability and number of offspring produced by the fM population at Day 3 improved threefold (3×) over the measurements for Generations 20 and 24 (Fig. 8). The egg hatchability (Fig. 8C) and the complete loss of fitness, or, conversely, the percentage of females remaining fertile as a function of age declined (Fig. 8D), and both of these measures showed accelerated declines in the fM when compared to the control. Unfortunately, the fitness measurement analyses of males from the (Fm) population were discontinued after Day 9 for COVID-19-related reasons, although a trend similar to that of the fM group was observed (Fig. 9). Both egg hatchability and the percentage of males remaining fertile decreased on an accelerated basis as a function of age (Fig. 9D). Taken together, both experimental populations (fM and Fm) showed a faster decline in the fertility of the younger sex under asymmetric mating.

**Fig. 8.**
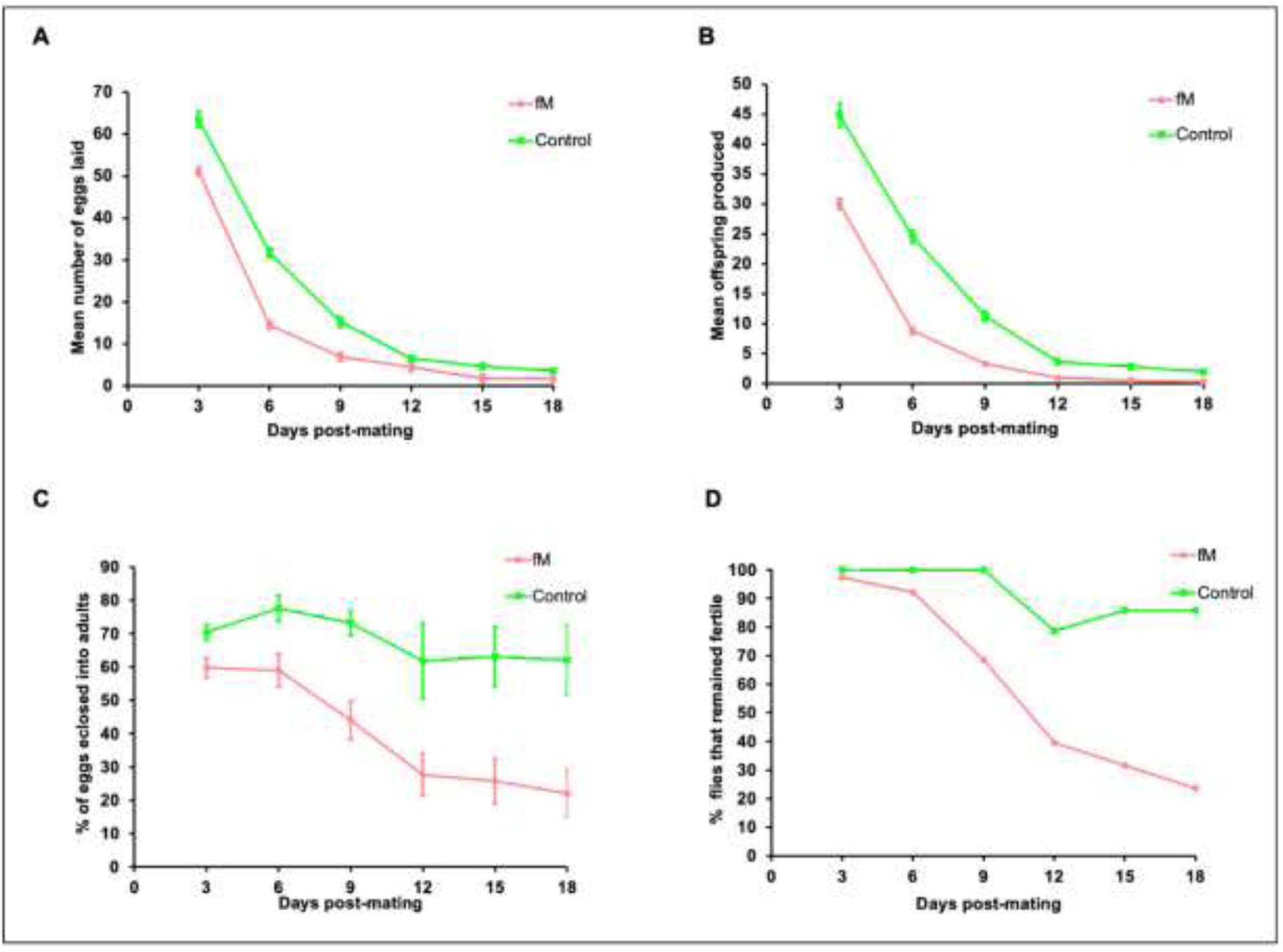
Fecundity and fertility curves for the control and young-female experimental (fM) populations. (A) The mean number of eggs laid per day, (B) mean number of offspring produced per day, (C) percentage of eggs eclosed per day, and (D) percentage of flies remaining fertile as a function of age, were measured every third day in the control and fM lines. Bars represent the standard error of the mean.

**Fig. 9.**
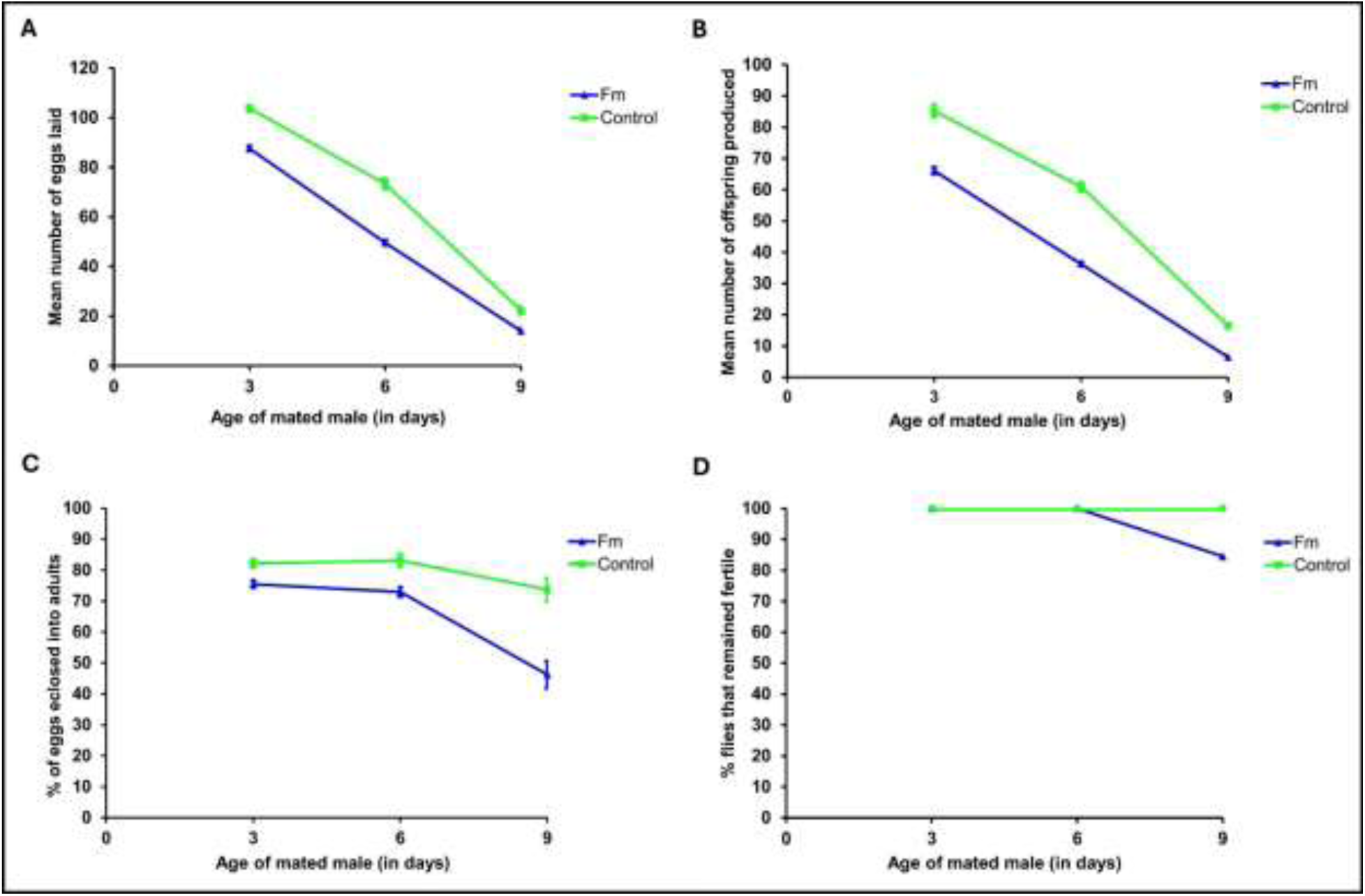
Fecundity and fertility curves for the control and young-male (fM) experimental populations. (A) The mean number of eggs laid per day, (B) mean number of offspring produced per day, (C) percentage of eggs eclosed per day, and (D) percentage of flies remaining fertile as a function of age, were measured every third day in the control and fM lines. Bars represent the standard error of the mean.

Our Weibull models used data from experimental population fM of Generation 70 (Fig 9 C) to demonstrate that the alternative models were superior to the null models at predicting the proportion of eggs eclosed (RSS null = 2753.61, RSS alternative = 33.51; p < 0.001; Fig.10 A & B) and the percentage of flies remaining fertile (RSS null = 4706.9, RSS alternative = 34.1; p < 0.001; Fig.10 C & D). These results indicate that the data are more accurately described when considering the control and experimental groups distinctly as opposed to combining the data without unique parameters. Therefore, the eclosion rate and number of flies remaining fertile declined faster in the experimental population than in the control.

**Fig. 10.**
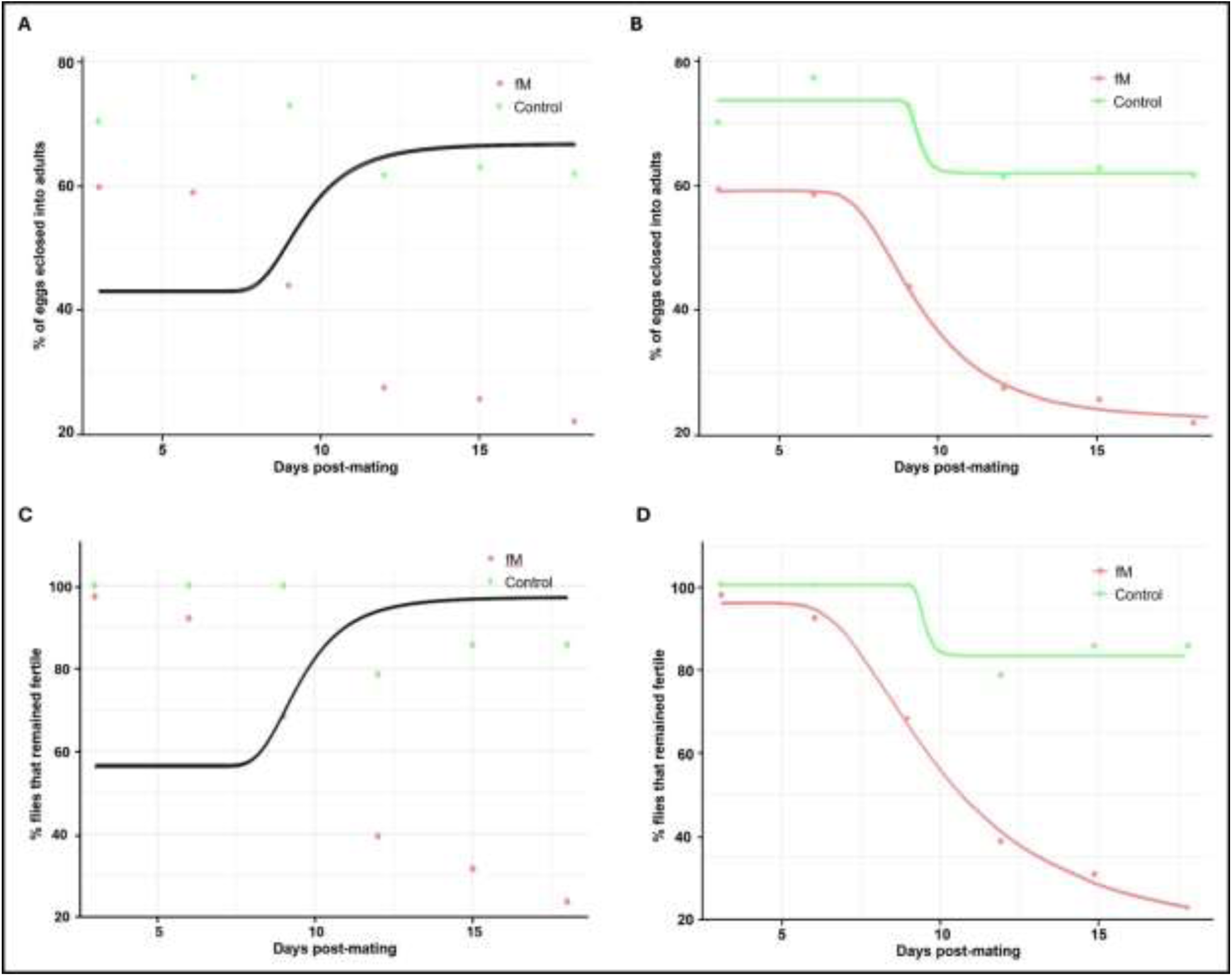
(A) the null model predicting the proportion of eggs that hatched over time, including both the control and experimental groups together in Generation 70. (B) the alternative model predicting the proportion of eggs that hatched over time, where the control group (in orange) and experimental group (in cyan) are each represented with independent parameters. (C) the null model predicting the proportion of flies remaining fertile over time, including both the control and experimental groups together. (D) the alternative model predicting the proportion of flies remaining fertile over time, where the control group and experimental group are each represented with independent parameters (based on graphs shown in **Fig. 8**).

Although the tests of fertility in experimental population Fm were discontinued after Day 9 and the data are insufficient for proper statistical analyses, the trend indicated a faster decline in eclosion rate and number of flies remaining fertile in the experimental line when compared to the control (Fig. 9 C & D).

## Discussion

The mate choice theory of menopause is not unusual in its operation, although it has a peculiar effect: It allows deleterious mutations to evolve as neutral mutations. Mate choice theory is consistent with our knowledge of the human mating system; namely, that throughout evolutionary history and in most global societies, a tendency exists for males to prefer younger mates (e.g., evolution from promiscuity to polygamy to monogamy, serial monogamy, widowers remarrying, harem, or sexual relationships outside marriage). Unlike other theories of menopause, mate choice theory lends itself to the genetic investigative work performed in this study.

### Age-restricted asymmetric mating: expectations

This experiment focused on three factors that affect fertility, one of which was known and two were unknown. The known factor was the accumulation of mutations, while the unknown components were the developmental immaturity of mates and inbreeding. In this study, 3-day-old females were allowed to mate and lay eggs for three days, and then discarded using standard Drosophila experimental protocol. Our expectation was that continued reproduction using asymmetric mating would lead to the accumulation of infertility mutations as neutral mutations in the younger sex with no post-reproduction lifespan. These mutations would be expressed if the younger sex lived and reproduced after Day 3. The fertility of the older mate would not be affected beyond the normal effects of aging. While beneficial mutations may be rare [38], deleterious mutations are common, with those affecting fertility more prevalent than those involving viability. Hybrid sterility evolves faster than hybrid inviability in *Drosophila* species [39], with many loci affecting male sterility [40].

The first unknown factor was how imposing asymmetric mating involving young mates would affect reproduction. However, our decision to use 3-day-old flies as mates was reasonable because flies of this age can mate and reproduce and sufficient time was available (approximately two weeks) to observe patterns of (declining) fecundity and fertility before the flies became infertile due to aging. A downside of this approach was that the 3-day-old flies may not have been fully mature. However, this study used reciprocal experiments involving both sexes as younger mates; therefore, the effects of uncontrolled variables, such as standing variation and developmental immaturity, should have been distinguishable from those arising from infertility mutations.

The second unknown element affecting fertility was the possible effect of inbreeding. We used a moderate sample size of 60 (30 males and 30 females) for each replicate, as the loss of genetic variation was irrelevant due to the experiment focusing on mutation accumulation as opposed to selection. Although repeated asymmetric mating over a long period would lead to loss of variation and inbreeding, the latter should have affected all populations similarly, as similar sample sizes were used in all populations and replicates.

### Expected and unexpected effects of asymmetric mating: observations

The fitness measurements taken from Generations 20, 24, and 70, showed some negative effects of asymmetric mating on reproductive capabilities on Day 3 (Fig. 1A-B, Fig. 2A-B, and Fig. 8A-B). Both the fM and Fm populations had reduced numbers of eggs laid, reduced hatchability of the eggs, and lower numbers of offspring produced at Day 3 when compared to the control. However, after Day 3, the hatchability of the eggs and the number of flies remaining fertile declined rapidly in Generations 20, 24, and 70 of both experimental populations. These results were interpreted as relating to the accumulation of deleterious mutations affecting fertility.

The female fitness reductions in Generations 20 and 24 were supported by declines in the number of ovarioles and stage-14 egg chambers. Ovariole numbers impact the production of eggs and the reproductive success of the flies [41, 42], and stage-14 denotes the mature primary oocyte that is ready for fertilization [43, 44], which is not produced until the female is 48 hours old [45]. Fewer ovarioles and stage-14 egg chambers were found in the fM population when compared to those observed in the Fm group and the control (Fig. 4).

While mate choice theory makes no predictions about lifespan, menopause discussions frequently link the evolution of fertility and longevity to senescence; therefore, we added this component to our investigation [25]. No significant difference was observed between the lifespans of the experimental and control populations, although a significant difference was noted between the lifespans of the males and females, whereby female flies lived longer than males (53 days vs. 33 days). These results support previous observations showing that virgin *Drosophila* females live longer than virgin males [46].

A significant difference in testis width was observed between the fM and Fm populations, but none was revealed between the fM group and the control (Fig. 6). Males from the Fm group had lower mean testis widths than males from the fM population during the early (age 0 to 15 days) and late (age 35 to 50 days) stages of the lifespan. As maturing cells move through the testis from the apical end to the seminal vesicle, spermatocytes become more predominant in the seminal vesicle and less evident in the testis [47]. The division of spermatocytes at the apical end of the testis slows with age [48], thereby reducing sperm production, while the movement of sperm toward the seminal vesicle continues. While this may account for the testis narrowing with age [47], the faster rate of decline in testis width during the later period (Days 25-50) suggests the effect of accumulated mutations. Mutation rates are higher during spermatogenesis than oogenesis [49].

### Mate choice theory is sex-neutral: both sexes can undergo menopause

The fecundity and fertility measurements in Generation 70 showed marked declines in the experimental populations as compared to the control, which corroborates the mate choice theory. In seeking to interpret these results, we should first elucidate the likelihood of observing the effects of such deleterious mutations. The mutation rates and their fitness effects have been investigated in previous population genetics and mutation accumulation studies (reviewed in [50]). The classic work of Terumi Mukai on mutation accumulations in *Drosophila* showed genome-wide mutation rates of 0.35-047 and a viability decline of approximately 1% per generation over 25-40 generations [51,52]. Subsequent studies using mutation accumulation and protein-coding sequences with different organisms have found mutation rates that vary by several orders of magnitude [50]; however, the *Drosophila* data on viability are of interest to us as benchmark measures for comparison with our fertility results.

Genes affecting fertility evolve faster than those affecting viability. Specifically, sex- and reproduction-related traits and genes evolve faster than non-sex traits and genes [53,54], hybrid sterility evolves faster than hybrid inviability [39], genes affecting male fertility evolve faster than genes affecting female fertility [55], and X-linked genes evolve faster than autosomal genes [56]. Furthermore, mutation rates are higher in males than females [49,57]. In general, sexual-system genes are quicker to evolve than genes affecting viability [58]; therefore, we expected mutation accumulation to affect fecundity and fertility in Generation 70 and our results confirmed this assumption. Despite the initial effects of using immature (3-day-old) mates and any impacts related to potential inbreeding, the accelerated fertility decline (egg hatchability) and complete loss of fertility as a function of age in both the fM and Fm populations verified the predictions of the mate choice theory. The decline of fertility in both experimental populations showed that menopause is not a peculiarity of females but of the sex reproducing at a younger age, whether male or female.

An important point to add here is that the rate of fitness decay in the two experimental populations, fM, and Fm, in generation 20 where we have full set of data for both populations, look rather similar. In view of the fact that rate of mutation is supposed to be higher in males than females, one may expect a faster fitness decay in the Fm population. This was not the case. This may be due to the fact that accumulation of deleterious fertility mutations by generation 20 may have been large enough to swamp any expected difference in the outcome of fM and Fm.

### Causes of menopause: A critique of the grandmother hypothesis

The limits of the grandmother hypothesis can be summarized as follows: Under the assumptions of the Red Queen hypothesis (the idea that species must constantly evolve to keep pace with the changing environment) grandmothers would have to “run” twice as fast as mothers to stay in the same place. This underlines the key weakness of the grandmother hypothesis [25]: The gain in fitness through kin selection is proportional to genetic relatedness, which declines by 50% in each filial generation. While kin selection can increase inclusive fitness, recuperating lost fitness through grandmothering is comparable to climbing a “steep fitness hill.” This is because a grandmother shares 50% of her genes with her children, but only 25% with her grandchildren.

This means that for every child that a grandmother did not bear, she would be required to raise two grandchildren over and above those produced by her children. Furthermore, for the grandmother strategy to evolve, the recuperation of lost fitness should occur immediately after menopause and not calculated over a future “prolonged lifespan” because evolution occurs based on current fitness, not potential future fitness.

The “live long” hypothesis [6], which is a sub-form of the grandmother hypothesis, explains the evolution of menopause in toothed whales based on the prolongation of the lifespan of grandmothers to overlap with their grandchildren. Infertility and longevity have separate genetic causes [59], and longevity cannot evolve through non-reproducing post-menopausal females [60] because, to reiterate the information in the prior paragraph, evolution occurs based on current fitness, not on the promise of future fitness.

Since the evolutionary oddity of menopause was first contemplated [3,31], the evolution of menopause and lifespan have been treated together because continued reproduction requires a continuation of lifespan and vice-versa. However, until recently [12], the potential independent evolution of the post-menopausal lifespan of females from the evolution of menopause had not been considered. A study of guppies (*Poecilia reticulata*) [60] reported that a significant proportion of individuals had post-reproductive lifespans that were unrelated to the length of their reproductive lifespans, which presented an interesting case of the evolution of a post-reproductive lifespan without menopause. Animals with short lifespans and annual reproduction can have post-reproductive lifespans because selection may be relaxed during the end part of their life. In contrast, long-lived animals rarely have post-reproductive lifespans without menopause. The rarity of menopause may imply that the evolutionary mechanism(s) involved are uncommon; therefore, age-based asymmetric mating may be an uncommon process.

## Conclusion

The mate choice theory predicts that a shift in mating behavior whereby older males prefer younger mates would lead to the accumulation of sex-specific, fertility-diminishing mutations that cause menopause in older females [12]. Age-restricted asymmetric mating with *D. melanogaster* resulted in a significant reduction in the fecundity and fertility of the younger mate, irrespective of sex. These results support the mate choice theory and show that changes in behavior, such as preferential mating, can have a profound impact on the reproduction and health of the female partners.

Mate choice theory is unique because it is a population-genetics-based theory that involves individual selection but excludes the founder effect, group selection, and kin selection; it operates by rendering deleterious mutations to evolve as neutral alleles, thereby avoiding the scrutiny of selection. In addition, it is symmetrical because it works irrespective of sex and either sex can become menopausal depending on which is used as the younger mate; this mate is consequently deprived of reproduction in old age. Menopause is rare, which may relate to the uncommon occurrence of the evolutionary mechanism(s) involved; therefore, age-based asymmetric mating may also be uncommon.

## Materials and Methods

### Rationale behind the study

The rationale behind this study was the desire to determine whether mating populations of young females only until a specific age (X) over many generations would result in the elimination of deleterious mutations affecting fertility until age X and the accumulation of neutral mutations after this age, which would affect fertility if the females were allowed to reproduce beyond age X. The theory is based on the observation that human males generally prefer younger mates, which deprives older females of reproduction [12]. Although human populations have no fixed or sharp cut-off point at which reproduction ends and male preference tapers off with advancing female age, for ease of experimentation with *Drosophila melanogaster* we used a cut-off point of 3-day-old females who were mated for 3 days.

### Base population

The flies used in this study came from a long-held laboratory base population derived from ten globally distributed *Drosophila melanogaster* strains that were sourced from the *Drosophila* Stock Center in California. The base population was maintained in the laboratory as a large population [61]. All experiments were conducted in laboratory conditions at room temperature (22°C) with flies reared on standard cornmeal-agar-molasses medium.

### Experimental protocol

Mate choice theory limits the age of the female (younger), while the males can be of any age. The theory can be evaluated by mating young females *en masse* with a mix of males of all ages to save time and labor. However, this method would have allowed for genetic variation and unintended selection; therefore, we attempted to reduce this possibility by mating 3-day-old young females (or males) with 20-day-old mature males (or females).

A mutation accumulation (for deleterious mutations affecting fertility) experiment was conducted using two experimental populations with an altered mating system compared to that of the random-mating control. The two experimental populations were created by using age-restricted, asymmetric mating with 3-day-old virgin females and 20-day-old males (fM), and 20-day-old females with 3-day-old virgin males (Fm). Three replicates were performed on all control and experimental populations.

*Drosophila melanogaster* females and males remained virgin for 8–10 hours and 12 hours after eclosion, respectively [62]. On Day 9, the jars containing the population of younger females mated with older males (fM) were cleared of adult flies when the pupae were ready to eclose, as shown in Fig. 11. On Day 10, 100 virgin males were collected, and 10 were placed in each food-filled vial. Males were left to age for 20 days and transferred into new vials every 7 days to prevent food contamination. On Day 26, the jars were cleared again, and 100 virgin females were collected on Day 27. On Day 30, the 20-day-old virgin males (30 individuals) were mated with the 3-day-old virgin females (30 individuals) in 3 jars (to avoid the risk of losing the line), and the progeny were used as the next generation. Given the 3-day mating window, when a sufficient supply of eggs had been laid on the food, the jars were cleared to start the new population, with the last day of mating becoming Day 0 for the next generation. Three independent replicates of the fM population were denoted as 1f, 2f, and 3f. The same procedure was used to create the older female × younger male population, denoted as Fm, with three independent replicates labeled as 1m, 2m, and 3m.

**Fig. 11.**
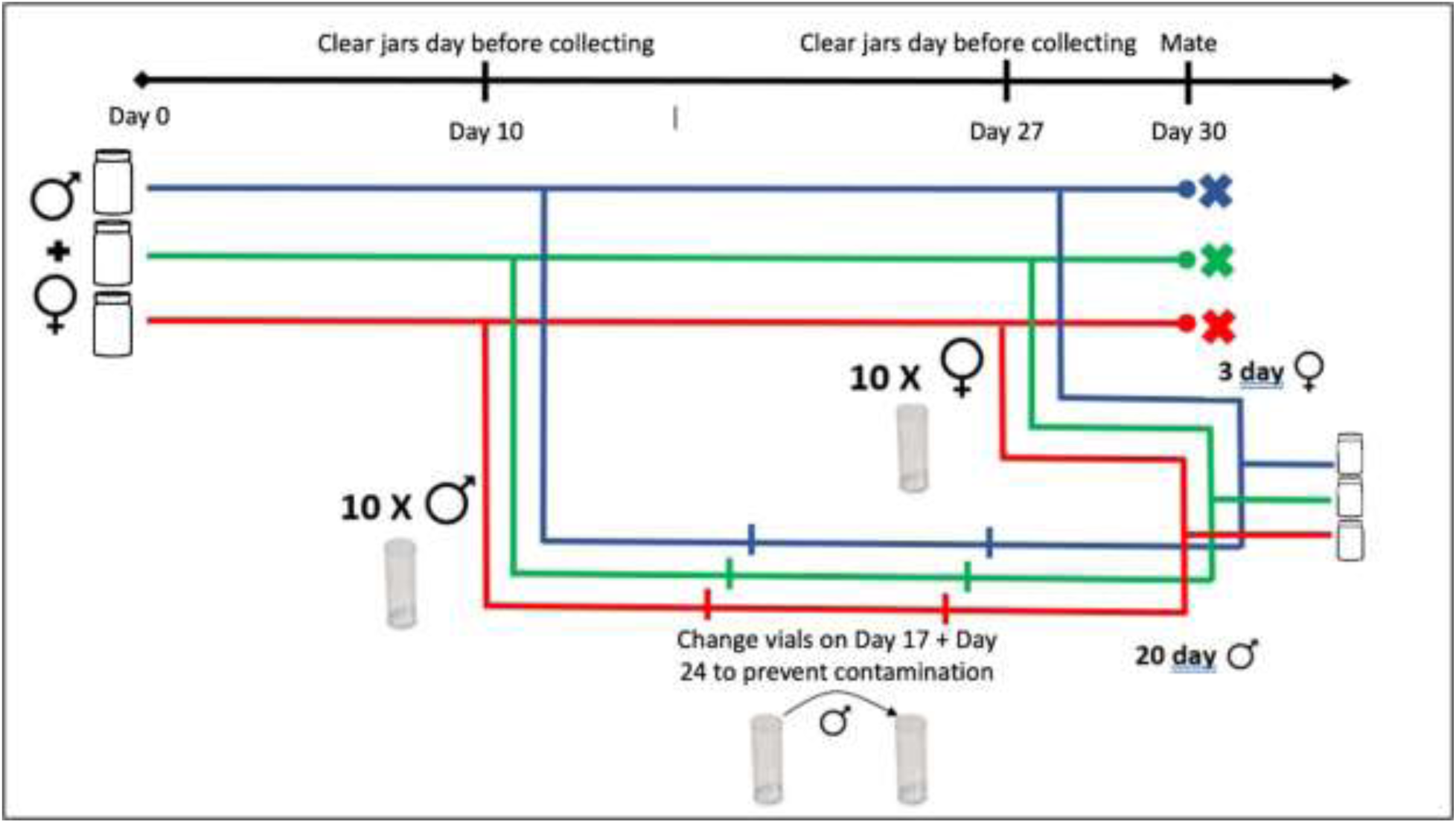
The age-restricted mating scheme used in the experimental populations of younger females mated with older males of *Drosophila melanogaster*.

### Measurement of fecundity and fertility

We measured four fitness components: the number of eggs produced, percentage of eggs eclosed, number of offspring produced, and percentage of post-mated flies remaining fertile. These measurements were made with flies from Generations 20, 24, and 70, after which the experiment was discontinued due to the COVID-19 pandemic.

For the fitness measurements, virgin individuals were sexed immediately after eclosion using CO2 and kept individually in food vials for 20 days. Subsequently, a 20-day-old male and a 3-day-old female from the same line were placed together in a vial and allowed to mate for three days, after which the male was removed and the female was allowed to oviposit. Females were transferred into a fresh vial every 24 hours to prevent larval overcrowding. The number of eggs laid during each 24-hour period was counted and used as a measure of fecundity, while the number of offspring eclosed was tallied every third day. Fertility was calculated as the proportion of eggs produced and eclosed (hatched) into viable offspring daily. The same procedure was concurrently conducted using 20-day-old females and 3-day-old males. After 3 days of mating, the male was removed and the female was allowed to oviposit.

Two controls were used during the fitness tests. For Generations 20 and 70, fitness measurements were made for both experimental lines, fM and Fm, and flies from the laboratory control population (a diversified base population) were used as experimental controls. For the experimental-line fitness measurements, the mating scheme used was the same as that used in the maintenance of the lines (fM, Fm). For the control fitness measurement, virgin males and females were kept individually in separate food vials and aged for 20 days, after which they were mated for a period of 3 days. On the third day, males were removed, females were allowed to oviposit, and fecundity and fertility were measured as described above. The mated flies from both the experimental and control groups came from within the line.

For the measurements using Generation 24, only one experimental population was examined (fM), *and the control, older males from the laboratory control population rather than from the experimental population (fM).* This arrangement differed from that of Generations 20 and 70, where the males in the experimental control came from within the population (fM) as explained in the previous paragraph. This was done to provide a male from the same population to see if it would make any difference. Since the mate choice theory depends on the age of the younger parent, the choice of the control male did not affect the outcome of the experiment. All tallies were made within a 20-day observation period that began after the 3-day mating period. A Student’s *t*-test or analysis of variance (ANOVA) was used to test for significant differences among means within and across populations (α = 0.05).

### Measurement of lifespan

Survivorship was measured using 50 males and 50 females for each of the three replicates of the experimental and the control populations (fM, Fm, control). Food jars (250 mL) were used, and all flies were virgins. The number of flies that remained alive was counted every third day, after which the surviving flies were transferred into fresh food jars. R software for Windows (Version 1.1.456) was employed to produce survival curves using the Kaplan–Meier survival method [36], and log-rank analysis [37] and Chi-square tests were performed with R software to compare populations.

### Measurement of ovariole and stage-14 egg chamber numbers

Virgin females from the experimental and control populations were collected post-eclosion and the number of ovarioles and stage-14 egg chambers were counted. Every third day, three females from each experimental (Fm and fM) and control replicate were sacrificed on a concave slide to observe the number of ovarioles and stage-14 egg chambers using a dissecting stereomicroscope. Each female was dissected into a drop of phosphate buffer solution (PBS) under the dissecting stereomicroscope. To identify and count individual ovarioles, 1 μL of crystal violet stain was added to each sample and fine needles were used to separate the eggs from the ovary. The number of eggs with dorsal appendages (stage-14 egg chambers) was counted from both the left and the right ovaries and recorded. The average count of the two ovaries was used as the stage-14 egg count per female.

The ovariole and stage-14 egg chamber numbers were obtained from 200 female flies, and the measurement was performed for 30 days post-eclosion. One-way ANOVA and two-tailed *t*-tests were performed to compare the populations using PRISM 6.0 software.

### Measurement of testis size

Testis size (length and width) was measured using an ocular micrometer under a compound light microscope. The ocular micrometer was calibrated using a stage micrometer to obtain the measurements in millimeters. Using a dissection microscope under 2× magnification, testes were dissected in PBS on a concave slide using forceps to pull them from the lower body. Individual testes were isolated and placed on glass slides. Each testis was stretched using fine forceps to straighten the coiling, viewed under a compound light microscope with 10× magnification, and measured by aligning the end of the testis with the ocular micrometer. The width of the testis at the midpoint was determined by dividing the length by 2 and measuring the width at the calculated length.

### Data analysis for egg hatching rate and flies remaining fertile

We aimed to determine the difference in the hatching rate and number of flies remaining fertile (or loss of fertility) between the experimental and control groups. For each of these assessments, we created a null model where both the control and experimental data were predicted by the same parametrization and an alternative model where the experimental and control groups were described by individual parameters.

To analyze the hatching rate (fly eclosion/eggs laid) and the number of flies that remained fertile each day, we used a four-parameter Weibull model for both factors:

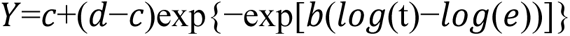

Here, *y* is either the proportion of eggs that hatched or the proportion of flies that remained fertile and *t* is time (in days). The null models considered one set of parameters for the control and experimental groups while the alternative models considered two distinct sets of parameters for groups. To determine which model had the better fit, we used an ANOVA and compared the RSS values.

## Acknowledgement

This research was supported by grants from the Natural Sciences and Research Council of Canada (NSERC) and the Faculty of Science, McMaster University, to RSS.

